# Detection of pathogenic *Leptospira* in the environment and its association with antagonistic *Pseudomonas* spp. and rainy season

**DOI:** 10.1101/2020.05.03.074963

**Authors:** K. Vinod Kumar, Chandan Lall, R. Vimal Raj, K. Vedhagiri, Anwesh Maile, N. Muruganandam, I. P. Sunish, Paluru Vijayachari

**Affiliations:** Regional Medical Research Centre (ICMR), WHO Collaborating Centre for Diagnosis, Reference, Research and Training in Leptospirosis, Port Blair-744101, Andaman and Nicobar Islands, India; Indian Council of Agricultural Research, National Institute of Veterinary Epidemiology and Disease informatics (ICAR-NIVEDI), Yelahanka, Bengaluru-560064, Karnataka, India; DPT - cGMP Facility, Central Research Institute, Ministry of Health & Family Welfare Kasauli. Himachal Pradesh - 173 204, India; ICMR-Regional Medical Research Centre, Bioinformatics division, Port Blair-744101, Andaman and Nicobar Islands, India

**Keywords:** Pathogenic *Leptospira*, Negative interaction, *Pseudomonas* spp., Surface water

## Abstract

Typically, humans contract leptospirosis through exposure to soil or water contaminated with the urine of infected animals. Specifically, people working in inundated fields, engaged in aquatic sports, or exposed to contaminated floodwater after periods of heavy rainfall bear the risk of contracting leptospirosis. There is a critical gap in the knowledge of the environmental cycle, transmission, and interaction of *Leptospira* species with its environment. A few studies establish the presence of higher concentration of leptospires during the rainy season when compared to the dry season. Therefore, we assessed the abundance of leptospires during the dry and wet months and their interaction with other microbes. The overall detection rate of leptospires in paddy field for the test period was 52 (49.5%). Leptospiral concentration positively correlated with the amount of rainfall (mm) during the sampling when compared to months that received comparatively less rainfall (60% vs. 28.5%, respectively). When observed for the microbial interaction, *Leptospira* showed significant negative correlation with *Pseudomonas* and rainfall in the paddy field. Moreover, Pseudomonas negatively correlated with the amount of rainfall. Corroborative results of *in-vitro* studies suggest the antagonistic effect of *Pseudomonas* spp. on leptospires. The results indicate that seasonal changes influence the diversity of free-living well-adaptive aquatic antagonistic microbe populations and may in turn determine the survival of *Leptospira* in the environment. Thus, microbial interaction can be the possible enigma for the fluctuation of *Leptospira* count in rainy and dry seasons in environmental surface water, which needs to be further confirmed. This will pave way for a better understanding of the survival of leptospires and the seasonal trend of exposure to humans.

## Introduction

Leptospirosis, a zoonotic disease having global prevalence, is caused by the pathogenic spirochete bacteria belonging to the genus *Leptospira*. Globally, around 1.03 million cases and 58,900 deaths due to leptospirosis are reported every year, which makes it a leading zoonotic cause of morbidity and mortality (Costa et al. 2015). Infection occurs when the susceptible host is exposed to bodily fluids (urine, placenta, and vaginal fluids) of an infected animal. Infection might also occur indirectly through exposure to contaminated moist soil and water – the most common cause of *Leptospira* infection.

Agricultural workers, especially those working in inundated fields, are at a higher risk of being infected with leptospires due to their day-to-day field-related activities (Tangkanakul et al. 2000). A long association of leptospirosis with paddy cultivation has been reported in tropical developing countries and is referred to as ‘rice field fever’ in many parts of the world (Antony 1996; Lall et al. 2016). Importantly, when the pathogen is shed into the environment and dispersed during heavy rains and flood, it can cause non-occupational exposure in urban and rural settings and can place the entire population at risk for leptospirosis. Leptospires survive in warm moist soil and water for several weeks and months (Levett 2001). Weather events, such as rain and flood, increase the risk of spreading infection. However, very few studies explain their presence during these events and information explaining how and why these factors influence the presence of *Leptospira* is missing, thereby rendering its control as a challenging task (Barragan et al. 2017). Therefore, a need to understand the environmental cycling, transmission, and interaction of *Leptospira* species with its environment is crucial.

In an aquatic environment, microbial communities depend on a complex interactive network of commensalism, antagonism, and parasitism. These biotic interactions are essential determinants of the natural microbial communities (Hibbing et al. 2010). A few studies have indicated that co-existing microbes might support *Leptospira* survival outside the host (Barragan et al. 2011; Kumar et al. 2015). Leptospires are known for biofilm formation with environment microbes as a protective mechanism, which may help in preventing dispersion in aquatic systems (Kumar et al. 2015; Kumar et al. 2016). Some microbes are competitive and actively eliminate the invading pathogens (Feichtmayer et al. 2017). It is difficult to predict the nature of pathogenic *Leptospira* in the environment, whether as an invading species dispensed in the environment by the reservoir/carrier host through urine or an autochthonous microorganism, having its habitat in water/soil and having a mechanism to survive like saprophytes.

*Pseudomonas fluorescens* is one of the most competitive, antagonistic, dominant, and well adapting group of bacteria found in diverse environments, especially agriculture fields, and produce a variety of compounds that facilitate their persistence in such habitats (Hibbing et al. 2010; Numberger et al. 2019). Although reports on effect of *Pseudomonas* on human pathogen in water/soil environment are scarce, its activity and mechanism against numerous plant pathogens and soil microbes are well established. There is no information on the vulnerability of leptospires to soil microbial shifts and on the presence of any specific microorganisms that can affect the survival of *Leptospira* in the environment in different climatic conditions. Thus, the present study was designed to test the hypothesis that the occurrence and persistence of pathogenic microorganism, such as *Leptospira*, are minimal in biologically active environments, such as water/soil, with dominant autochthonous bacteria, such as *Pseudomonas*, when compared to the sterile one as described earlier (Feichtmayer et al. 2017). With respect to the influence of climatic factors, the interaction between the two bacteria was investigated under rainy and non-rainy seasons.

## Materials and Methods

### Water sample collection

The Andaman and Nicobar Islands of India are located in a subtropical region, situated between 6°N and 14°N latitude and 92°E and 94°E longitude, and are known to be endemic for leptospirosis. The water samples were collected from paddy fields of four villages, which were reported to be endemic pockets for leptospirosis in South Andaman district: Manglutan, Wandoor, Chidiyatapu, and Chouldari. The rainfall data for the study period was accessed from the Indian Meteorological Department, Government of India website (http://www.imd.gov.in/section/hydro/distrainfall/webrain/bayislands-/andaman.txt.) and http://andssw1.and.nic.in/ecostat/2012/rainfall.pdf

### In-situ q-PCR quantification of Leptospira and Pseudomonas spp

Water/soil samples (15 v/ml each) were collected in sterile polypropylene centrifuge tubes from paddy fields (n=105). The samples were centrifuged at 6,000 × g for 20 minutes. DNA was extracted from the sediment by using QIAamp DNA Mini Kit (Qiagen, California, US) following the manufacturer’s protocol. The quantitative real-time PCR assay (SYBR Green method) was performed targeting the *Pseudomonas*-specific and *Leptospira*-specific primers, viz., Pseu16S: Forward -5’ CTA CGG GAG GCA GCA GTG G -3’ and Reverse -5’ TCG GTA ACG TCA AAA CAG CAA AGT -3’ (Purohit *et al.,* 2003), and LipL 32: Forward 5’-ATC TCC GTT GCA CTC TTT GC-3’ and Reverse 5’-ACC ATC ATC ATC ATC GTC CA-3’(Ahmed et al. 2006; Kumar et al. 2015), respectively. These primers were earlier used for conventional PCR and thus optimized prior to the q-PCR SYBR Green assay using RT-PCR system 7500 (Applied Biosystems, CA, USA). The reaction mixture was prepared using the Power SYBR Green PCR Master mix 2X kits (Applied Biosystems, CA, USA), according to the manufacturer’s protocol. The temperature profile include two holding stages, *viz*., 2 min at 50°C and 2 min at 95°C, followed by 40 cycles of amplification (95°C for 25s, 55°C for 45s, 72°C for 45s) after which the reaction was stopped (96°C for 2 min) and melted (60&#8211;96°C with plate readings). Standard curves for quantification of *Leptospira interrogans*, strain Pomona and *P. putida* isolate R-1 were plotted (Ganoza et al. 2006). For preparing standards, leptospires and pseudomonads were counted using a Petroff Hauser counting chamber and were serially diluted with sterile double-distilled H2O to make 10^8^ to 10^0^ leptospires ml^−1^ and genomic DNA was prepared subsequently by using the QIAamp DNA Mini Kit. The experiment was performed in triplicates to generate the standard curve and obtain plain sterilized ultrapure water as a control. The test was considered negative when no amplification was yielded before the 40^th^ cycle.

### Isolation of *Pseudomonas*

Soil/water samples (500 gm or ml) were collected from paddy fields. Samples were transported in sterile polyethylene bags to the laboratory, serially diluted, and plated on *Pseudomonas* selective medium, King’s medium B (KMB) (King et al. 1954), supplemented with chloramphenicol (13 μg ml^−1^), cyclohexamide (100 μg ml^−1^) and ampicillin (50 μg ml^−1^) (Simon and Ridge 1974). After incubation of the plates at 28°C for 48 hours, the colonies were checked for fluorescence under UV light (Sharifi Tehrani et al. 1998) and fluorescent colonies were picked for further sub-culturing on fresh KMB agar plates to obtain pure cultures. The cultures were stored in stab agar overlaid with 20% glycerol stock, and kept at −80°C until further use.

### Identification of pseudomonads

The 17 purified cultures were subjected to DNA isolation using the methodology described by Ausubel et al. (1999). Polymerase chain reaction (PCR) was performed by using 16S rDNA primers: Forward -5’ AGA GTT TGA TCM TGG CTC AG -3’ and Reverse-5’ ACG GCT ACC TTG TTA CGA CTT -3’ with standard protocol (Lane et al. 1985). The amplified products were sequenced using DNA analyzer 3730 (Applied Biosystems, California, USA). All sequences were assembled using the software MEGA 5 (Tamura et al. 2011) and aligned using CLUSTALW with default parameters. A phylogenetic dendrogram was generated using neighbor-joining tree of MEGA 5.

### *In-vitro* effect of *Pseudomonas* spp. on the survival of leptospires

The inhibitory effect of *P. putida* isolate R-1 on leptospires was studied by co-culturing them in two different media, EMJH liquid medium (pH 7.4) and sterile paddy field water (pH 7.3). The paddy field surface water samples were collected (1.5 liters each) in sterile poly-vinyl containers from the villages of Manglutan and Chouldari in South Andaman district. The water samples were pooled, sterilized, and filtered using 0.22μm filter (MILLEX^®^GV). *P. putida* isolate R-1 cell suspensions were prepared in varying concentrations of approx. 1.6 – 2.2 × 10^1^, 1.6 – 2.2 × 10^2^, 1.6 – 2.2 × 10^3^, 1.6 – 2.2 × 10^4^, 1.6 – 2.2 × 10^5^, 1.6 – 2.2 × 10^6^ and 1.6 – 2.2 × 10^7^ cells ml^−1^ in the respective experimental conditions, i.e., in EMJH medium and paddy field surface water. Log phase cultures of each *Leptospira* strains in EMJH medium were centrifuged for 10 min at 5,000 × g, washed once with Phosphate Buffer Saline (PBS) pH 7.4, and re-suspended in the respective medium to a concentration of 2.8 × 10^7^ cells ml^−1^.

The co-culture mixture was prepared with 100 μl suspension for each strain of *L. interrogans* Pomona and *L. biflexa* Patoc 1. These were added separately to 9.9 ml of the *P. putida* isolate R-1 cell suspension, at a concentration of 2.8 × 10^5^ leptospiral cells ml^−1^. The mixture was incubated at room temperature (25-28°C) and the leptospiral count and motility was assessed through standard methods (i.e., average bacterial numbers in five cell counts) using Petroff Hauser counting chamber (Hauser Scientific, Hosham, Pennsylvania, US) and DFM (Zeiss AXIO, Germany) every 24 hours until the 40th day. Thereafter, the assessment was made every 20^th^ day until the 30^th^ day comparing with the test control.

The co-culture mixture was stained with live/dead BacLight Bacterial Viability Assay Kit (Invitrogen) for leptospiral viability and for visualization of the live and dead cells, by following the manufacturer’s instructions. Images were acquired by fluorescent microscopy (AXIO Scope A1, Zeiss, Germany). The leptospiral viability was also checked by filtering the mixture with 0.22 filters and inoculating the filtrate into a fresh EMJH semi-solid medium followed by incubation at 28±2°C. The culture tubes were checked for a period of three months using DFM for any leptospiral growth. All experiments were performed in triplicates.

### Identification of the active compounds by GC/MS

*P. putida* isolate R-1 was grown in 100 ml of kings B media. Culture supernatants were extracted twice with an equal volume of ethyl acetate. The mixture was concentrated, weighed, and suspended in 1 mg ml^−1^ concentration in 0.1% methanol. Minimal Inhibitory Concentration (MIC) of the extract for leptospiral strains was determined by microdilution method, as described for antibiotics (Murray and Hospenthal 2004). For gas chromatography/mass spectrometry (GC/MS), 10 mg of the extract was suspended in 1 ml ethyl acetate Agilent GC-MS-5975C instrument (Santa Clara, California) equipped with a column DB-5ms and was used for the analysis. The GC oven was held at 130°C for 2 min and was then increased to 300°C for 8 min. Helium was used as the carrier gas with a flow of 30 ml min-1. The mass spectra results were analyzed in NIST-05 software. The identification of compounds was based on 90% similarity between the MS spectra of unknown and reference compounds in an MS spectra library.

### Statistical analysis

The results were analyzed and represented by using GraphPad Prism 5.04 (San Diego, California, United States) and SPSS v16.0 for Windows 7. The bacterial genome count for the two organisms in the environmental water samples for the studied season was compared using paired sample ‘*t*’ test and Pearson’s correlation coefficient, correlation analysis and Past program (Hammer et al. 2001). The effect of *Pseudomonas* on leptospires concentration *in vitro* was assessed by survival analysis and plot for respective medium by using Kaplan-Meier analysis in SPSS v16.0 software and by normalizing the value to log.

## Results and Discussion

### Quantification and correlation of leptospires and *Pseudomonas* in the environment

Totally, 105 samples were quantified for pseudomonads and leptospires by q-PCR, and peaks less than the threshold value were considered as zero. The overall detection rate of the test period was 52 (49.5%) positives, which ranged between 0 and 15745 (n=105) cells ml^−1^ for leptospires [mean, 3047.15 cells ml^−1^ (95% confidence interval, 2073.97-4020.34)]. The detection rate of leptospires varies with the study’s geographical location. The detection rate ranged from 15.9% to 33.3% in waterlogged paddy field soil in an earlier study conducted in Andaman (Kumar et al. 2016; Lall et al. 2016). Other high rates of detection around the world include 56.69% from soil samples in New Caledonia (Thibeaux et al. 2018), 48% in Philippines (Sato et al. 2019), 33.3% in water from Malaysia (Kelantan) (Mohd Ali et al. 2018), 34% in standing water in Brazil (Salvado, Pau da Lima) (Casanovas-Massana et al. 2018a) and 30.6 % (33/108) in soil in Taiwan (Fuh et al. 2011). However, a few studies report the quantification of *Leptospira* in environmental samples like water and soil. Ganoza et al. (2006) reported mean concentrations around 1,000 cells/mL in urban area waters, Casanovas-Massana et al. (2018a) reported 152 to 166 GEq/mL, and Vein et al. (2012) reported a concentration of 10^4^ genome-equivalents/mL in environmental water samples. A recent study by Casanovas-Massana et al. (2018b) has indicated that *Leptospira* can be present in the soil and water at concentrations below the detection limit of PCR and qPCR. Thus, the concentration of bacteria in samples may be much more than what is described in the present study and elsewhere. Whereas on the contrary, molecular techniques may overestimate the presence of organisms, as PCR is based on DNA that can be from both viable and non-viable dead *Leptospira* cells (Casanovas-Massana et al. 2018b).

Enough evidence suggests the association of human and animal leptospirosis outbreak and upsurge with high rainfall, which is considered as a primary risk factor for leptospirosis worldwide (Barragan et al. 2017; Sato et al. 2019). However, there are few studies establishing the presence and concentration of causative organisms (leptospires) during the rainy season. In the present study, the overall leptospiral concentration in paddy field positively correlated with the amount of rainfall (mm) during the sampling (*r=0.541, d.f.=105, P<0.001*). The results were also analyzed by distributing the observation into rainy and non-rainy months, i.e., June to November 2011 (n=35), December 2011 to May 2012 (n=35), and June to November 2012 (n=35), with a mean rainfall of 422.36 (±211.97 SD), 142.77 (±143.89 SD), and 411.56 (±236.05 SD). The detection rates of *Leptospira* were higher in samples collected during the rainy months than the months that received comparatively less rainfall (60% vs. 28.5%, respectively). The concentration of leptospires collected during the rainy months (June to November) significantly correlated (*r = 0.533, d.f.= 35, P = 0.001*) with the amount of rainfall. Leptospires have been reported in water and soil samples after periods of heavy rainfall and even during summer (Barragan et al. 2017). A similar trend was observed in Brazil where the detection of leptospires was higher in sewage samples collected during the rainy season (47.2%) than the dry season (12.5%) (Casanovas-Massana et al. 2018a). A study using next generation sequencing found that the number of leptospiral sequences significantly correlated with the amount of rainfall (mm) on the sampling day, even after two days after the rainfall. Surprisingly, the study reported that other suspected environmental parameters, such as temperature, humidity, and weather, had no correlation with the concentration of leptospires at the time of collection (Sato et al. 2019).

These studies provide evidence that there is an apparently high concentration of leptospires during the rainy season. However, as reported by Barragan et al. (2017), we can only speculate about how conditions during rain or floods favor the increased leptospires’ concentration in the environment, even though abiotic factors like physiological parameters, pH, temperature, humidity, rainfall, and soil nutrients have been studied (Lall et al. 2018). Until the present, the exploration of biotic factors is limited to biofilm formation and coexistence of beneficial microbes. Bacteria associated with leptospires include *Sphingomonas* spp. (Barragan et al. 2011), *Azospirillum brasilense* (Kumar et al. 2015), and proteobacteria (α-, β-, γ-, and δ-proteobacteria); and one species each of *Actinobacteria*, *Flavobacteria*, and intracellular bacteria *Chlamydiae* (Sato et al. 2019). Although Sato et al. (2019) studied the entire diversity of 16s RNA sequences in environmental samples using DNA meta-barcoding, they have only presented the correlation of associated bacteria and did not mention anything about the negatively correlated microbes in the sample that were devoid of *Leptospira* DNA. Moreover, the variability of *Leptospira* concentration across the season has never been attributed to the antagonistic microbes.

The present study explores the presence of antagonist bacterial species and a possible mechanism that could influence the *Leptospira* concentration in rainy and non-rainy seasons. There can be a number of antagonist microbes in such environments. However, this study focused on one antagonistic microbe *Pseudomonas* spp. and the rationale for choosing this bacterium may be debatable, as there are no previous reports on the effect of pseudomonads on *Leptospira* or any other spirochetes. It has been chosen for its merits of abundance in cultivation field and reputation of antagonistic nature toward many bacteria, especially in agricultural fields. In the present study, when same water samples (n=105) were quantified for pseudomonads by q-PCR, the overall detection rate of the test period was 74 (70.5%) positives that ranged between 0 and 18971 cells ml^−1^ [mean, 3004.52 cells ml^−1^ (95% confidence interval, 2095.07-3913.97)]. The pair wise test of *Leptospira* conc. *vs. Pseudomonas* conc. showed a significant negative correlation *(r=− 0.320, d.f.=105, P=0.001*) in paddy field. Among the 105 samples tested, *Pseudomonas* was detected in 38 samples (36.19%) and *Leptospira* was detected in 16 samples (15.23%). Both organisms were observed in 36 samples (34.28 %), while 15 samples (14.28%) did not detect any of these organisms. On the other hand, the concentration of pseudomonads in paddy field negatively correlated with the amount of rainfall (*r=− 0.419, d.f.=105, P<0.001*) (Fig. 1). When distributed in rainy and non-rainy months, the concentration of leptospires positively correlated with the amount of rainfall, whereas concentration of pseudomonads significantly correlated (*r=0.238, d.f.=105, P<0.001*) with non-rainy months (December to May) (Fig. 1, Supplementary Table S1, Fig. S1). Similarly, lower abundance of *Pseudomonas* and other microbial diversity in the rainy season was reported by other researchers (Rodriguez-Verdugo et al. 2012; Smit et al. 2001). Thus, a reduction in antagonistic agent in the ecosystem may facilitate the proliferation of *Leptospira* during the rainy season.

**Figure 1.**
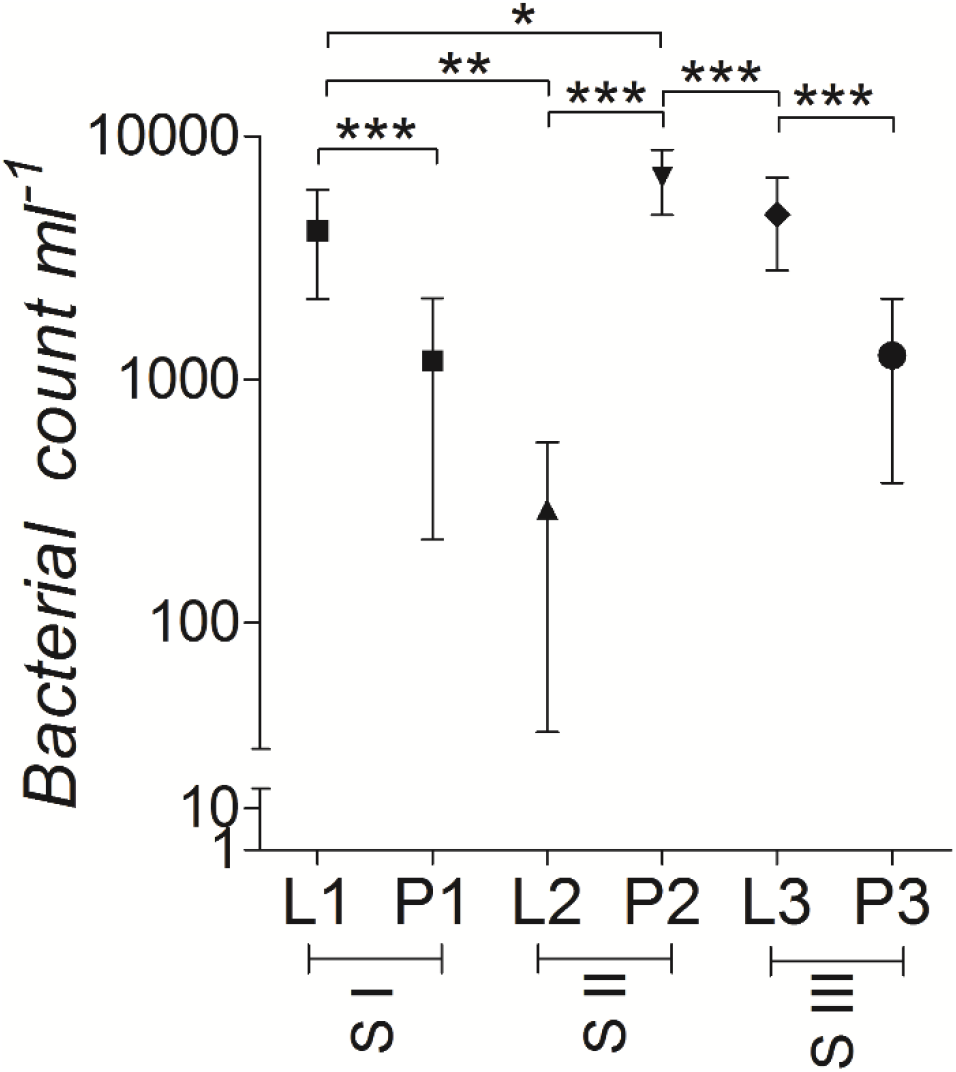
Quantification of *Leptospira* and *Pseudomonas* in paddy fields, S I (June to November 2011) Versus S II (December 2011-May 2012) Versus S III (June 2012-Nov 2012). Bars indicate 95% confidence intervals. Analysis of variance, F= 11.4, *p<* 0.0001. Newman-Keuls multiple comparison test *p*-value represented as *p*>0.0005 (***), *p*>0.005 (**), *p*>0.05 (*).

Leptospires are not so fragile spirochetes, as earlier studies have shown that they can survive and maintain virulence in unfavorable conditions in water for a long time (Barragan et al. 2017). Paddy field seems to provide a favorable condition for the growth and survival of leptospires and *Pseudomonas*, as very few paddy fields were found devoid of these bacteria in this study. This study would be important to understand the mechanism of increase in *Leptospira* concentration in environmental surface waters, especially during rainfall and floods. The limitation of this study would be the studying of interaction between two species in an environment with diverse microbial flora. However, one single or few species can sometimes act as keystone species, which control the existence of other microbes in the environment (Hibbing et al. 2010). Environmental *Pseudomonas* spp., especially *Pseudomonas fluorescens* group, would be the ideal antagonistic microbes to study against *Leptospira* because of its abundance and dominance in agricultural fields. These bacteria thrive in diverse conditions and niche of water and soil ecosystems, are better adapted, and produce diverse compounds to compete other microbes that facilitate their persistence in such habitats when compared to any other bacteria (Hibbing et al. 2010; Numberger et al. 2019).

### Diversity of *Pseudomonas* in the paddy field

Totally, 17 fluorescent *Pseudomonas* isolates were recovered from the soil/water. The phylogenetic analysis suggested that all 17 environmental isolates belonged to *P. fluorescens* lineage (Table S2 supplementary material). The isolate R1, used for *in-vitro* studies, belonged to *P. putida* group (Acc. No. JX112041.1) and other paddy field isolates belonged to three different groups. Among the total isolates analyzed, 82.3% (14) of the isolates belonged to *P. putida* group; 11.76% (2) belonged to *P. fluorescens* group, and 5.88% (1) belonged to *P. aeruginosa* group (Fig. 2). These three groups are reported to be dominant antagonistic agents and their application is found in various fields (Sarniguet et al. 1995). Although a negative association was demonstrated between *Pseudomonas* spp. and *Leptospira* community, it was not possible to investigate the impact of individual species of *Pseudomonas* or *Leptospira*, due to their mixed nature and extreme species diversity in the field condition by PCR. Besides, our study focused mainly on the diversity and influence of *Pseudomonas* spp. on the *Leptospira* community *In-situ,* while the *in-vitro* mechanisms of such interactions are still unknown.

**Figure 2.**
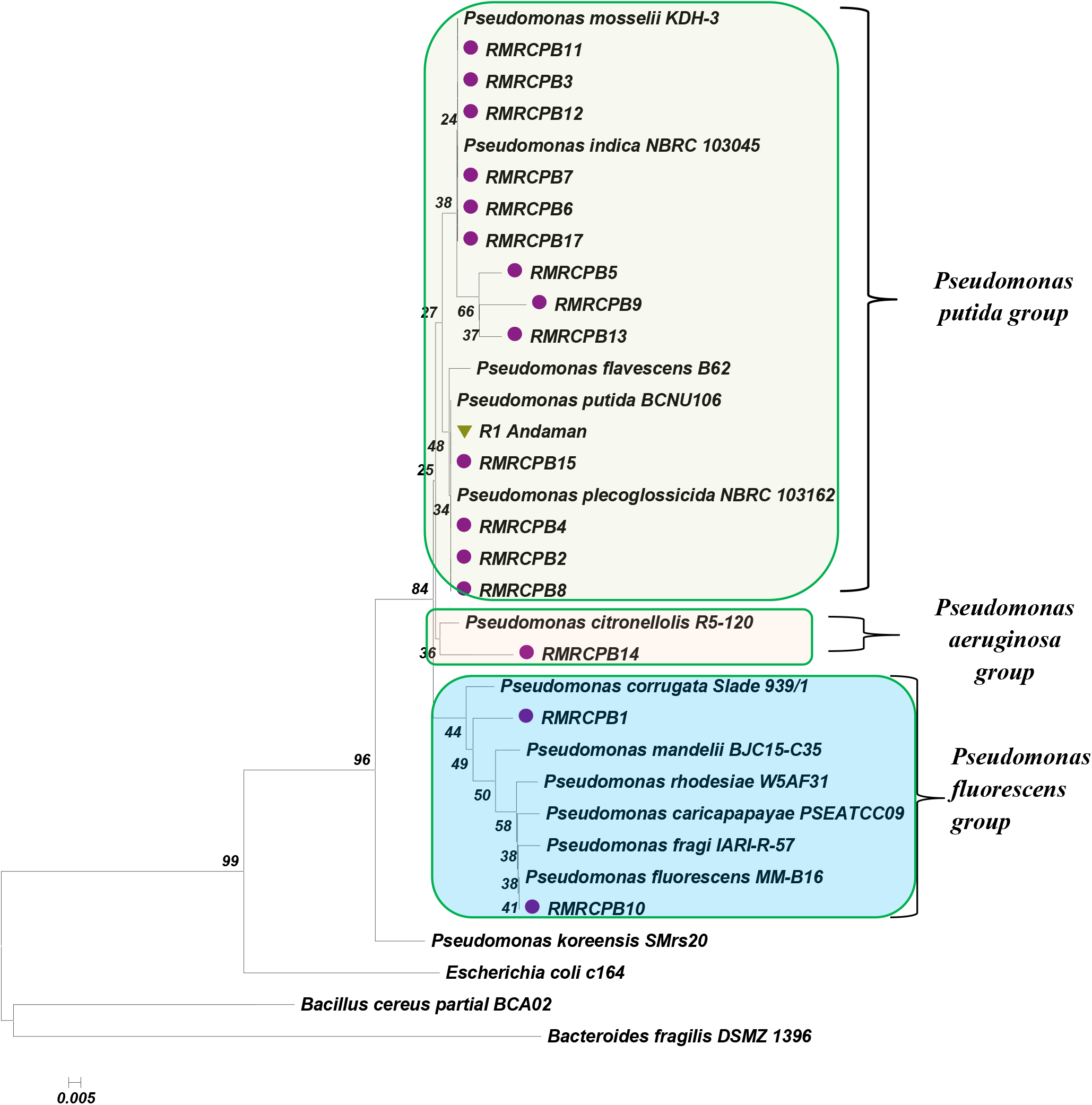
The inferred phylogenetic tree of environmental fluorescent *Pseudomonas* including 17 fluorescent *Pseudomonas* isolates and 16 reference strains. Evolutionary distances were determined with pairwise dissimilarities of the 16S rRNA gene sequences, and the dendrogram was generated using the neighbor-joining algorithm (Mega 5). Here (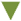 green triangle) isolate R1 Andaman used for *In-vitro* testing against leptospires, (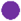 purple circle) Environmental *Pseudomonas* isolates for the determination of the environmental diversity of pseudomonads.

### Effect of Pseudomonas on the survival of Leptospires in a dual species microcosm

In order to further prove this phenomenon of negative interaction, an attempt was made to create a dual species microcosm for *Leptospira* and *Pseudomonas* tested *in-vitro.* Thus, one of the isolates, *P. putida* R-1, was selected for testing against *Leptospira* and was found effective in inhibiting the leptospires at a dose of 1.6 × 10^2^ cells ml^−1^ in both EMJH and paddy field water. The live/dead BacLight Bacterial Viability Assay showed the death of leptospires cells when cultured with *P. putida* isolate R-1 (Fig. 1 a1, a2). The survival curves for the reduction of leptospires’ count in the respective medium was created by Kaplan-Meier analysis, as shown in Fig. 3 b1, b2 and Supplementary Table S3, S4. The presence of *P. putida* isolate R-1 in the EMJH medium was lethal to the growth of leptospires and the count declined completely in three days for pathogenic (mean 1.50, CI 95% – 0.24, 2.77) and for saprophytic (mean 1.00, CI 95% −0, 2.13). In paddy field surface water, the leptospiral counts were reduced to 100% within 8 to 10 days for *L. interrogans* (mean-3.63, CI 95% – 1.78, 5.48) and for *L. biflexa* (mean 5.91, CI 95% – 3.20, 8.62), when treated with *P. putida* isolate R-1 and compared to control (Supplementary Table S5). The average cell count of leptospires (leptospires alone) in paddy field water (under controlled condition) was approx. 5.0 × 10^5^ cells ml^−1^ (mean-8.17 × 10^4^, CI-95% 2.6 × 10^4^ – 1.37 × 10^5^) on the 30^th^ day (*t=*20.74*, d.f.=15, P<*0.001), which was higher than the initial concentration and indicated growth in paddy field water. *Leptospiral* strains used in the present study underwent multiple *in-vitro* passages; however, these strains adapted well to the paddy field surface water and might have contained stimulatory factors for their survival (Picardeau et al. 2008). This hypothesis is supported by previous studies, which reported that *L. interrogans* had genetic machineries for living a dual life, viz., both in the mammalian host and in surface water, while *L. biflexa* had a well-established genetic foundation for environmental survival (Picardeau et al. 2008; Ren et al. 2003).

**Figure 3.**
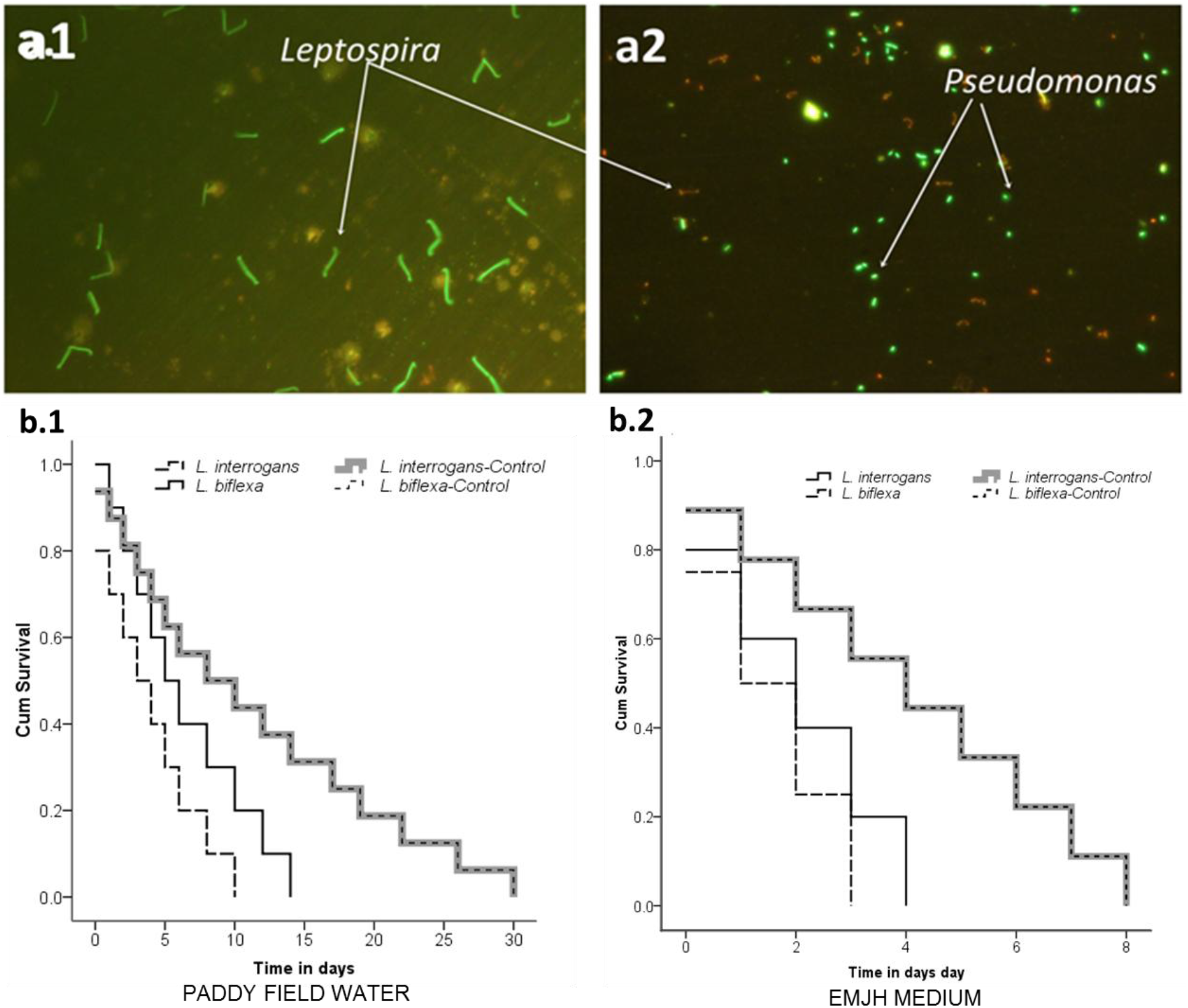
*In-vitro* analysis of *Leptospira* strains co-cultured with *Pseudomonas* spp. R-1 in Paddy Field surface water and in EMJH medium. a) Leptospires co-cultured with pseudomonads and stained with Live/Dead BacLight Bacterial Viability kit, a1) Leptospires alone (Control) green fluorescence under fluorescence microscope (40×) indicates viable cells and a2) Green rods indicates viable Pseudomonads and orange indicates dead leptospires. Survival curves of *L. interrogans* and *L. biflexa* treated with *P. putida* R1 in b.1) paddy field water and b.2) EMJH medium constructed by Kaplan-Meier analysis

A similar deleterious effect of environmental isolate *Flavobacterium* spp. and *Delftia* spp. on *Leptospira in-vitro* was reported by Barragan et al. (2011). The study also shows that not all environmental bacteria exert inhibitory effect on leptospires. *Sphingomonas* spp. exerted a positive effect on leptospires in distilled water at a concentration range of 1.9 × 10^6^ cells/ml to approx. 4.9 × 10^7^ cells/ml (Barragan et al. 2011). In our earlier study, two bacteria, viz., *Azospirillum* sp. (Kumar et al. 2015) and *Azotobacter* spp. were observed to demonstrate a positive effect on the survival and growth of leptospires.

It is not clear how the interaction between *Pseudomonas* and *Leptospira* may work in a diverse environment. The presence of other beneficiary microbes may influence the interaction between these two species, which needs to be further investigated. The deleterious effect of *Pseudomonas* may be attributed to the fact that it is a highly antagonistic organism. In the present study, it was also found that this particular isolate, belonging to the group of plant growth promoting bacteria (Supplementary Table S3), is antagonistic by nature to most of the plant pathogens. Moreover, *Pseudomonas* is a promising bio-control agent against many plant fungal pathogens, viz., *Rastonia solanacearum, Fusarium oxysporum Rhizoctonia solani, Septoria tritici, Thielaviopsis basicola, Pythium ultimum*, etc. (Gao et al. 2012; Pal et al. 2001) and bacterial pathogens, viz., *Xanthomonas oryzae, Ralstonia solanacearum, Xanthomonas campestrisb,* etc. (Mondal et al. 2001; Wang et al. 2005). However, their impacts on human pathogenic organisms are still unknown.

The antagonistic effects of *Pseudomonas* on plant pathogens are nutrient depletion, Hydrogen cyanide (HCN) production, bioactive compound production, and the action of siderophore (high-affinity ferric chelators). These effects deplete the iron content in the environment. Iron is the growth-limiting factor for both plant pathogens (Compant et al. 2005) and for the pathogenic *Leptospira* (Asuthkar et al. 2007). In contrast, it has been reported that the leptospires could survive in a nutrient-depleted condition (Trueba et al. 2004) and studies argue that *Leptospira* could acquire iron from different sources, including siderophores of other bacteria, which is not investigated (Louvel et al. 2006). Bioactive compounds produced by these environmental strains could be responsible for their deleterious effect on leptospires.

Further, the MIC of the Ethyl acetate extract of the culture supernatant of the *P. putida* isolate R-1 inhibited leptospiral growth at a conc. of 50 μg ml^−1^ and the MBC was at a conc. of 100 μg ml^−1^. The GC-MS analysis of ethyl acetate extract of *P. putida* isolate R-1 indicated the presence of three compounds. The identification of active principles is based on their retention time (RT), molecular formula, molecular weight (MW), concentration (peak area %), and peak of the compounds (Supplementary Fig. S2). The major component identified was 2-Piperidinone, whereas, Benzene-propanol and Pyrrolo [1, 2-a] pyrazine-1, 4-dione derivatives were also identified in significant amount in the tested extract. 2-Piperidinone is known for its antimicrobial activity (Zhou et al. 2014).

## Conclusion

Outside the host (human/animal), *Leptospira* is known to have survival strategies like cell aggregation (Trueba et al. 2004) and biofilm formation (Ristow et al. 2008), which enable them to overcome abiotic stress in the environment. Interactions of *Leptospira* with other microbes in the environment are not much known. Hence, this work holds significance from the ecological point of view to understand the survival and transmission of *Leptospira* in the environment and outside the host. There are possibilities of two scenarios: *Leptospira*, as an invading species, can be dispensed in the environment by the reservoir/carrier host in the urine; and pathogenic *Leptospira*, as native species, can have the ability to survive like saprophyte in the environment. Laboratory results suggest the vulnerability of *Leptospira* in the presence of *Pseudomonas* spp. The fluctuation in the presence of antagonistic microbes may determine the survival of *Leptospira* in environment in either of the cases (Fig. 4). Thus, microbial interaction can be the possible answer for the enigma of fluctuation of *Leptospira* count in rainy and dry seasons in environmental surface water. However, it is premature to conclude that the single genus *Pseudomonas* controls the presence of leptospires in the environment. Thus, this will encourage further studies in the area of *Leptospira* microbial interaction.

**Figure 4.**
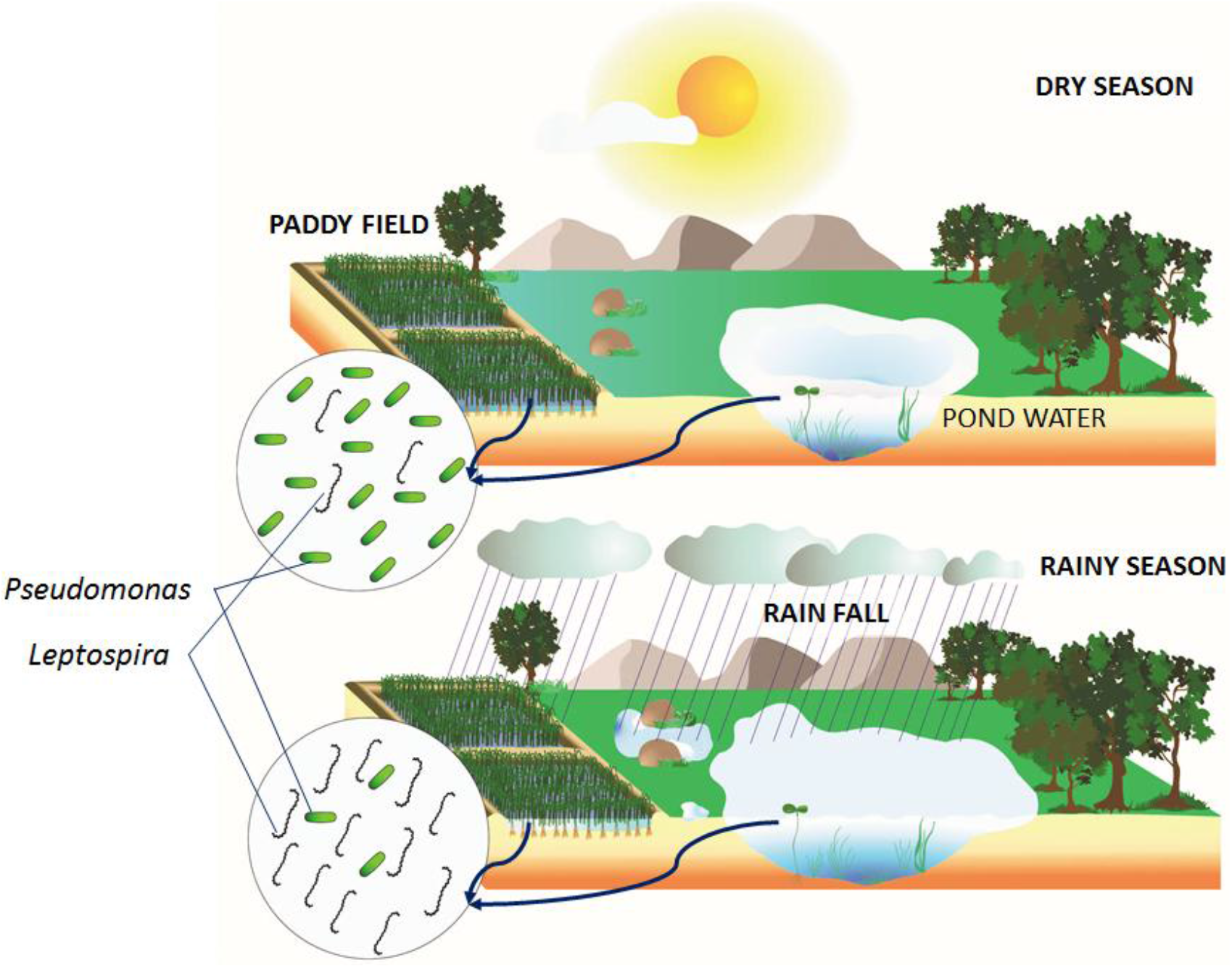
Illustration of effect of seasons on *Leptospira* and *Pseudomonas* interaction in environmental water body.

## Acknowledgement

The authors are thankful to the Indian Council of Medical Research, New Delhi for providing intramural grants for the study.

## Supplementary material

**Table S1:** correlation analysis of count of *Leptospira* and *Pseudomonas* within and among the wet and dry seasons

**Table S2**. Identification of the paddy field biofilm isolates by alignment with 16S rRNA gene sequences of published bacterial species in the NCBI database.

**Table S3:** Cumulative survival curve description for effect of *P. putida* R1 on leptospires growth and viability in EMJH medium

**Table S4:** Cumulative survival curve description for effect of *P. putida* R1 on leptospires growth and viability in water medium

**Table S5**. Mean and median period (in days) of *Leptospira* reduction when treated with *P. putida* R1 strain in the paddy field surface water and EMJH medium *in-vitro* condition.

**Figure S1**. Detection of *Leptospira* and *Pseudomonas* DNA in the environmental samples during wet and dry months. Comparative rate of detection for leptospires (dark blue horizontal bar) and *Pseudomonas* (brown horizontal bar), detection of different species of *Pseudomonas* (Yellow bar), average rain fall (dull blue).

**Figure S2.** GC-MS spectra of ethyl acetate extract of *Pseudomonas putida* R1 showing peaks

