## Supplemental material for "Detection of pathogenic *Leptospira* in the environment and its association with antagonistic *Pseudomonas* spp. and rainy season"

### Supplementary material

#### Supplementary tables

**Table S1: Correlation analysis of count of *Leptospira* and *Pseudomonas* within and among the wet and dry seasons**

| | Leptospires S1 | Pseudomonads S1 | Leptospires S2 | Pseudomonads S2 | Leptospires S3 | Pseudomonads S3 |
|---|---|---|---|---|---|---|
| **Leptospires S1** | 1 | -0.261 | -0.143 | 0.252 | 0.509** | 0.056 |
| **Pseudomonads S1** | -0.261 | 1 | -0.065 | -0.142 | -0.245 | -0.135 |
| **Leptospires S2** | -0.143 | -0.065 | 1 | -0.326 | -0.291 | -0.141 |
| **Pseudomonads S2** | 0.252 | -0.142 | -0.326 | 1 | 0.430** | -0.185 |
| **Leptospires S3** | 0.509** | -0.245 | -0.291 | 0.430** | 1 | -0.154 |
| **Pseudomonads S3** | 0.056 | -0.135 | -0.141 | -0.185 | -0.154 | 1 |

*Here S1: June –November 2011, S2: December 2011 to May 2012 and S3: June – November 2012, **. Correlation is significant at the 0.01 level (2-tailed). *. Correlation is significant at the 0.05 level (2-tailed).*

**Table S2. Identification of the paddy field biofilm isolates by alignment with 16S rRNA gene sequences of published bacterial species in the NCBI database.**

| Isolate | Sequence length (bp) | Closest published database accession | Proposed identity | % Identity |
|---|---|---|---|---|
| **R1 Andaman** | 740 | KJ819570.1 | *Pseudomonas putida* | 99 |
| **RMRCPB1** | 150 | KF135229.1 | *Pseudomonas* spp. | 99 |
| **RMRCPB2** | 150 | KP313546.1 | *Pseudomonas putida* | 99 |
| **RMRCPB3** | 150 | KF202609.1 | *Pseudomonas* spp. | 99 |
| **RMRCPB4** | 140 | KP313564.1 | *Pseudomonas putida* | 99 |
| **RMRCPB5** | 132 | KM242105.1 | *Pseudomonas aeruginosa* | 99 |
| **RMRCPB6** | 148 | KF202556.1 | *Pseudomonas* spp. | 99 |
| **RMRCPB7** | 131 | KF202556.1 | *Pseudomonas* spp. | 100 |
| **RMRCPB8** | 129 | KF313546.1 | *Pseudomonas putida* | 99 |
| **RMRCPB9** | 149 | HM56592.1 | *Pseudomonas* spp. | 99 |
| **RMRCPB10** | 136 | KP126778.1 | *Pseudomonas fluorescens* | 99 |
| **RMRCPB11** | 145 | KF202609.1 | *Pseudomonas* spp. | 100 |
| **RMRCPB12** | 133 | JF488194.1 | *Pseudomonas* spp. | 99 |
| **RMRCPB13** | 144 | JQ316364.1 | *Pseudomonas* spp. | 99 |
| **RMRCPB14** | 118 | KJ524111.1 | *Pseudomonas* spp. | 99 |
| **RMRCPB15** | 150 | KP313357.1 | *Pseudomonas putida* | 99 |
| **RMRCPB17** | 148 | JQ316364.1 | *Pseudomonas* spp. | 99 |

**Table S3: Cumulative survival curve description for effect of *P. putida R1* on leptospires growth and viability in EMJH medium**

| Test | | Time in days | Status | Cumulative Proportion Surviving at the Time | | N of Cumulative Events | N of Remaining Cases |
|---|---|---|---|---|---|---|---|
| | | | | Estimate | Std. Error | | |
| ***L. interrogans*** | 1 | 0 | 5.483 | 0.8 | 0.179 | 1 | 4 |
| | 2 | 1 | 5.472 | 0.6 | 0.219 | 2 | 3 |
| | 3 | 2 | 5.269 | 0.4 | 0.219 | 3 | 2 |
| | 4 | 3 | 4.529 | 0.2 | 0.179 | 4 | 1 |
| | 5 | 4 | 0 | 0 | 0 | 5 | 0 |
| ***L. biflexa*** | 1 | 0 | 5.458 | 0.75 | 0.217 | 1 | 3 |
| | 2 | 1 | 5.228 | 0.5 | 0.25 | 2 | 2 |
| | 3 | 2 | 5.131 | 0.25 | 0.217 | 3 | 1 |
| | 4 | 3 | 0 | 0 | 0 | 4 | 0 |
| ***L. interrogans-Control*** | 1 | 0 | 5.415 | 0.889 | 0.105 | 1 | 8 |
| | 2 | 1 | 5.301 | 0.778 | 0.139 | 2 | 7 |
| | 3 | 2 | 5.602 | 0.667 | 0.157 | 3 | 6 |
| | 4 | 3 | 6.699 | 0.556 | 0.166 | 4 | 5 |
| | 5 | 4 | 7.602 | 0.444 | 0.166 | 5 | 4 |
| | 6 | 5 | 7.954 | 0.333 | 0.157 | 6 | 3 |
| | 7 | 6 | 8.212 | 0.222 | 0.139 | 7 | 2 |
| | 8 | 7 | 8.301 | 0.111 | 0.105 | 8 | 1 |
| | 9 | 8 | 0 | 0 | 0 | 9 | 0 |
| ***L. biflexa-Control*** | 1 | 0 | 5.301 | 0.889 | 0.105 | 1 | 8 |
| | 2 | 1 | 5.602 | 0.778 | 0.139 | 2 | 7 |
| | 3 | 2 | 6.699 | 0.667 | 0.157 | 3 | 6 |
| | 4 | 3 | 7.602 | 0.556 | 0.166 | 4 | 5 |
| | 5 | 4 | 7.954 | 0.444 | 0.166 | 5 | 4 |
| | 6 | 5 | 8.212 | 0.333 | 0.157 | 6 | 3 |
| | 7 | 6 | 8.301 | 0.222 | 0.139 | 7 | 2 |
| | 8 | 7 | 8.342 | 0.111 | 0.105 | 8 | 1 |
| | 9 | 8 | 0 | 0 | 0 | 9 | 0 |

**Table S4: Cumulative survival curve description for effect of *P. putida* R1 on leptospires growth and viability in water medium**

| Test | | Time in days | Status | Cumulative Proportion Surviving at the Time Estimate | Cumulative Proportion Surviving at the Time Std. Error | N of Cumulative Events | N of Remaining Cases |
|---|---|---|---|---|---|---|---|
| | 1 | 0 | 5.471 | 0.889 | 0.105 | 1 | 8 |
| | 2 | 1 | 5.384 | 0.778 | 0.139 | 2 | 7 |
| | 3 | 2 | 5.342 | 0.667 | 0.157 | 3 | 6 |
| | 4 | 3 | 5.295 | 0.556 | 0.166 | 4 | 5 |
| ***L. interrogans*** | 5 | 4 | 5.166 | 0.444 | 0.166 | 5 | 4 |
| | 6 | 5 | 5.052 | 0.333 | 0.157 | 6 | 3 |
| | 7 | 6 | 4.865 | 0.222 | 0.139 | 7 | 2 |
| | 8 | 8 | 4.45 | 0.111 | 0.105 | 8 | 1 |
| | 9 | 10 | 0 | 0 | 0 | 9 | 0 |
| | 1 | 0 | 5.458 | 0.909 | 0.087 | 1 | 10 |
| | 2 | 1 | 5.432 | 0.818 | 0.116 | 2 | 9 |
| | 3 | 2 | 5.41 | 0.727 | 0.134 | 3 | 8 |
| | 4 | 3 | 5.374 | 0.636 | 0.145 | 4 | 7 |
| | 5 | 4 | 5.342 | 0.545 | 0.15 | 5 | 6 |
| ***L. biflexa*** | 6 | 5 | 5.269 | 0.455 | 0.15 | 6 | 5 |
| | 7 | 6 | 5.228 | 0.364 | 0.145 | 7 | 4 |
| | 8 | 8 | 5.073 | 0.273 | 0.134 | 8 | 3 |
| | 9 | 10 | 4.927 | 0.182 | 0.116 | 9 | 2 |
| | 10 | 12 | 4.705 | 0.091 | 0.087 | 10 | 1 |
| | 11 | 14 | 0 | 0 | 0 | 11 | 0 |
| | 1 | 0 | 5.8 | 0.938 | 0.061 | 1 | 15 |
| | 2 | 1 | 5.45 | 0.875 | 0.083 | 2 | 14 |
| | 3 | 2 | 5.45 | 0.812 | 0.098 | 3 | 13 |
| | 4 | 3 | 5.458 | 0.75 | 0.108 | 4 | 12 |
| | 5 | 4 | 5.473 | 0.688 | 0.116 | 5 | 11 |
| | 6 | 5 | 5.481 | 0.625 | 0.121 | 6 | 10 |
| | 7 | 6 | 5.506 | 0.562 | 0.124 | 7 | 9 |
| ***L. interrogans-*** | 8 | 8 | 5.528 | 0.5 | 0.125 | 8 | 8 |
| ***Control*** | 9 | 10 | 5.541 | 0.438 | 0.124 | 9 | 7 |
| | 10 | 12 | 5.561 | 0.375 | 0.121 | 10 | 6 |
| | 11 | 14 | 5.586 | 0.312 | 0.116 | 11 | 5 |
| | 12 | 17 | 5.597 | 0.25 | 0.108 | 12 | 4 |
| | 13 | 19 | 5.619 | 0.188 | 0.098 | 13 | 3 |
| | 14 | 22 | 5.629 | 0.125 | 0.083 | 14 | 2 |
| | 15 | 26 | 5.639 | 0.062 | 0.061 | 15 | 1 |
| | 16 | 30 | 5.643 | 0 | 0 | 16 | 0 |
| | 1 | 0 | 5.458 | 0.938 | 0.061 | 1 | 15 |
| | 2 | 1 | 5.441 | 0.875 | 0.083 | 2 | 14 |
| | 3 | 2 | 5.456 | 0.812 | 0.098 | 3 | 13 |
| | 4 | 3 | 5.471 | 0.75 | 0.108 | 4 | 12 |
| | 5 | 4 | 5.491 | 0.688 | 0.116 | 5 | 11 |
| | 6 | 5 | 5.522 | 0.625 | 0.121 | 6 | 10 |
| | 7 | 6 | 5.557 | 0.562 | 0.124 | 7 | 9 |
| ***L. biflexa-Control*** | 8 | 8 | 5.577 | 0.5 | 0.125 | 8 | 8 |
| | 9 | 10 | 5.577 | 0.438 | 0.124 | 9 | 7 |
| | 10 | 12 | 5.596 | 0.375 | 0.121 | 10 | 6 |
| | 11 | 14 | 5.608 | 0.312 | 0.116 | 11 | 5 |
| | 12 | 17 | 5.614 | 0.25 | 0.108 | 12 | 4 |
| | 13 | 19 | 5.643 | 0.188 | 0.098 | 13 | 3 |
| | 14 | 22 | 5.665 | 0.125 | 0.083 | 14 | 2 |
| | 15 | 26 | 5.685 | 0.062 | 0.061 | 15 | 1 |
| | 16 | 30 | 5.7 | 0 | 0 | 16 | 0 |

**Table S5. Mean and median period (in days) of Leptospira reduction when treated with *P. putida* R1 strain in the paddy field surface water and EMJH medium in vitro condition.**

| | Test | Mean | Std. Error | 95% Confidence Interval | Median | Std. Error | 95% Confidence Interval | Log Rank (Mantel-Cox) |
|---|---|---|---|---|---|---|---|---|
| In Paddy field water | *L. interrogans* | 4.33 | 1.09 | 2.19, 6.48 | 4 | 1.49 | 1.08, 6.92 | *$\chi^2$ =9.29, d.f. =3, P=0.026* |
| | *L. biflexa* | 5.91 | 1.39 | 3.2, 8.62 | 5 | 1.65 | 1.76, 8.24 | |
| | *L. interrogans-Control* | 11.19 | 2.33 | 6.62, 15.75 | 8 | 4 | 0.16, 15.84 | |
| | *L. biflexa-Control* | 11.19 | 2.33 | 6.62, 15.75 | 8 | 4 | 0.16, 15.84 | |
| | *Overall* | 8.89 | 1.13 | 6.68, 11.09 | 6 | 1.2 | 3.65, 8.35 | |
| In EMJH Medium | *L. interrogans* | 2 | 0.71 | 0.61, 3.39 | 2 | 1.1 | 0, 4.15 | *$\chi2$ = 6.85, d.f. =3, P=0.077* |
| | *L. biflexa* | 1.5 | 0.65 | 0.24, 2.77 | 1 | 1 | 0, 2.96 | |
| | *L. interrogans-Control* | 4 | 0.91 | 2.21, 5.79 | 4 | 1.49 | 1.08, 6.92 | |
| | *L. biflexa-Control* | 4 | 0.91 | 2.21, 5.79 | 4 | 1.49 | 1.08, 6.92 | |
| | *Overall* | 3.26 | 0.49 | 2.31, 4.21 | 3 | 0.64 | 1.75, 4.25 | |

#### Supplementary figures

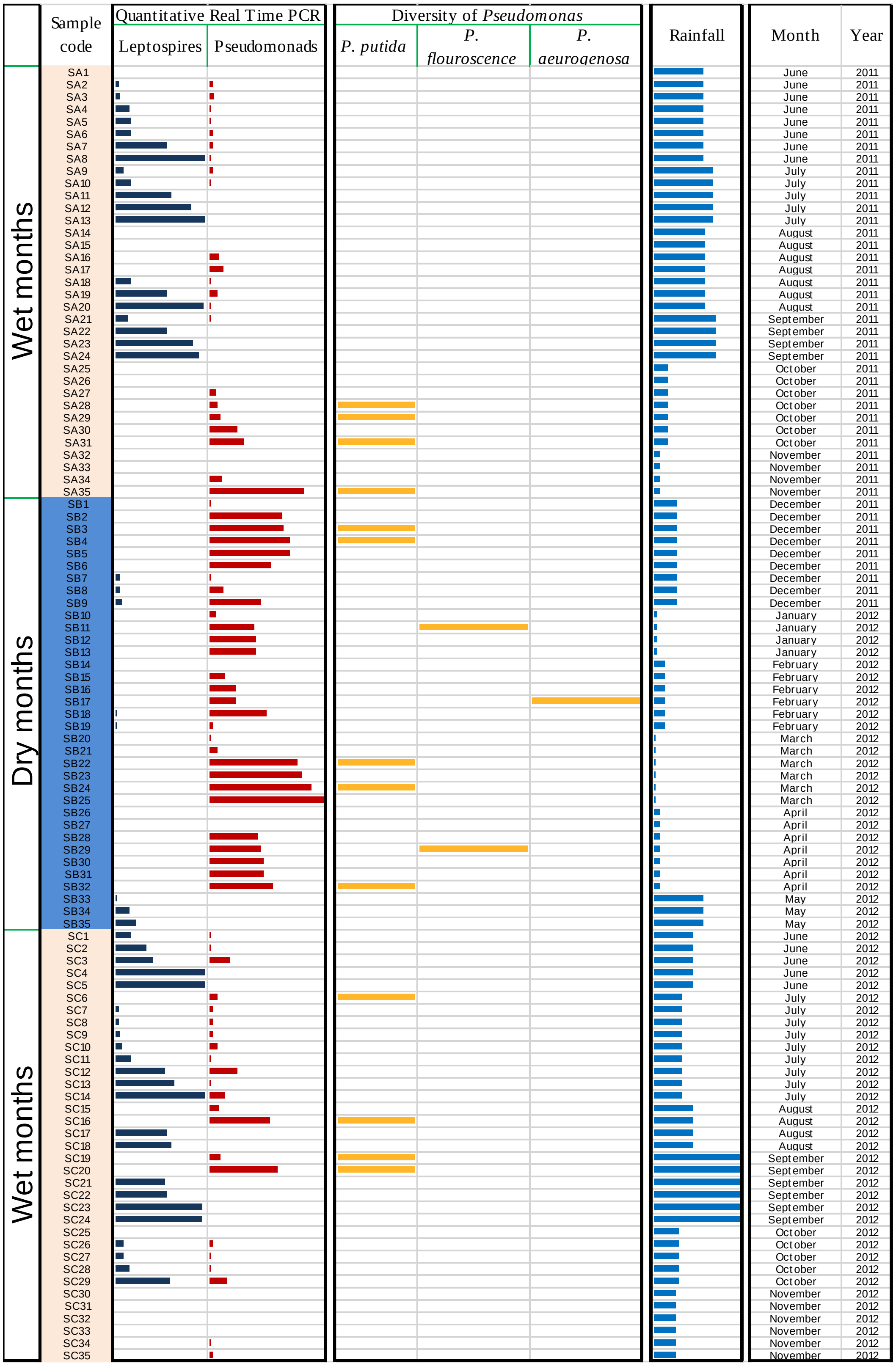


**Figure 1.** Detection of *Leptospira* and *Pseudomonas* DNA in the environmental samples during wet and dry months. Comparative rate of detection for leptospires (dark blue horizontal bar) and *Pseudomonas* (brown horizontal bar), detection of different species of *Pseudomonas* (Yellow bar), average rain fall (dull blue).

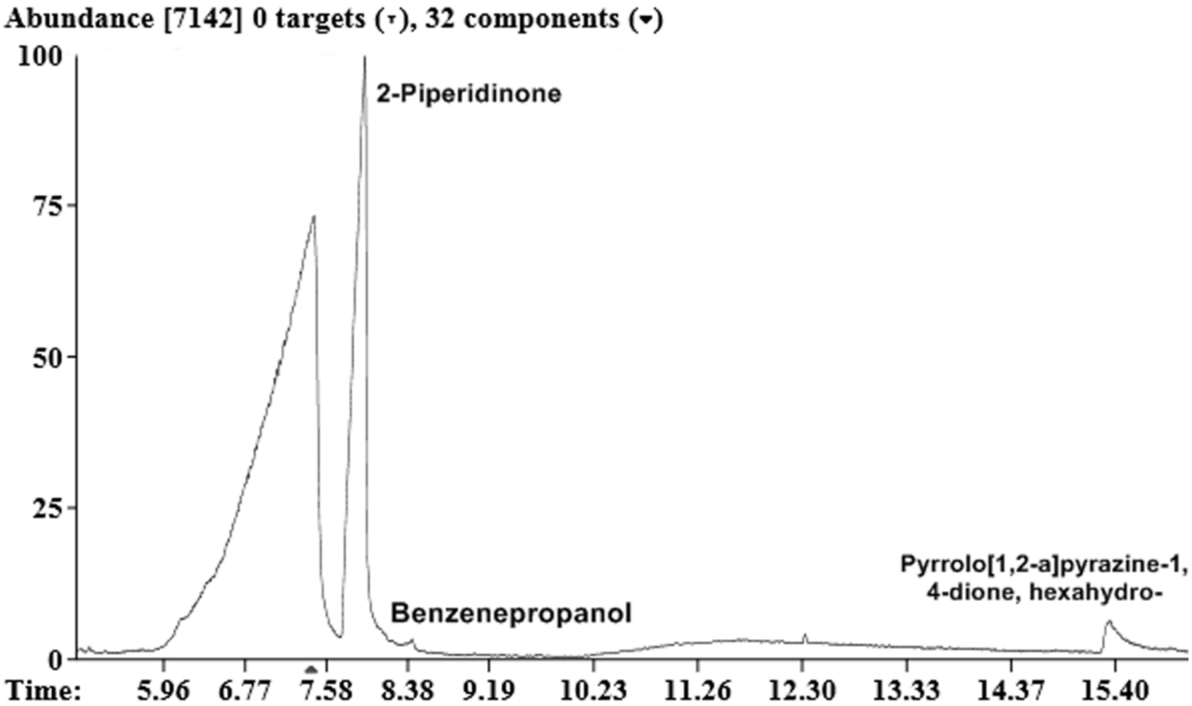


**Figure S2.** GC-MS spectra of ethyl acetate extract of *Pseudomonas* putida R1 showing peaks
